# Multifaceted contributions of *Dicer2* to arbovirus transmission by *Aedes aegypti*

**DOI:** 10.1101/2022.11.14.516413

**Authors:** Sarah Hélène Merkling, Anna Beth Crist, Annabelle Henrion-Lacritick, Lionel Frangeul, Valérie Gausson, Hervé Blanc, Artem Baidaliuk, Maria-Carla Saleh, Louis Lambrechts

## Abstract

Arthropod-borne viruses (arboviruses) transmitted by *Aedes aegypti* mosquitoes are an increasing threat to global health. The small interfering RNA (siRNA) pathway is considered the main antiviral immune pathway of insects, and thus represents a potential key target in the development of novel transmission-blocking strategies against arboviruses. Although the antiviral function of the siRNA pathway in *Ae. aegypti* is well established, its effective impact on arbovirus transmission is surprisingly poorly understood. Here, we used CRISPR/Cas9-mediated gene editing to create a line of *Ae. aegypti* in which we mutated *Dicer2*, a gene encoding the RNA sensor and key component of the siRNA pathway. Our *Dicer2* null mutant line is viable and fertile, and only displays minor fitness defects despite being unable to produce siRNAs. The loss of *Dicer2* affects early viral replication and systemic viral dissemination of four medically significant arboviruses (chikungunya, Mayaro, dengue, and Zika viruses) representing two viral families. However, measures of virus transmission potential indicate that *Dicer2* null mutants and wild-type mosquitoes display an overall similar level of vector competence. Additionally, *Dicer2* null mutants undergo significant virus-induced mortality during infection with chikungunya virus, but not dengue virus. Together, our results define a multifaceted role for *Dicer2* in the transmission of arboviruses by *Ae. aegypti* mosquitoes and pave the way for further mechanistic investigations.

## Introduction

Mosquitoes are vectors of several medically significant arthropod-borne viruses (arboviruses) that have been emerging or reemerging over the past decades^1^. Despite significant efforts to control mosquito populations, develop specific antiviral drugs, or deploy effective vaccines, arboviral diseases constitute a growing threat for global health^2^. The main arbovirus vectors are the mosquito species *Aedes aegypti* and *Aedes albopictus*, which are distributed extensively on all continents and extend their geographical range rapidly^3^. A mosquito acquires an arbovirus through ingestion of a blood meal taken from a viremic vertebrate host. In the digestive tract, the virus crosses the epithelial barrier and infects midgut cells^4^. After a few days, the virus exits the midgut and disseminates to the hemocoel, infecting other organs and cells, such as the fat body and hemocytes^4^. Eventually, the virus reaches the salivary glands, where it replicates^5^, before being retransmitted to another vertebrate host upon the next infectious bite^4^.

At every step of the mosquito infection, arboviruses encounter tissue barriers and innate immune responses. Major immune signaling pathways such as Toll, immune deficiency (IMD), and Janus kinase/signal transducers and activators of transcription (JAK-STAT) pathways have been implicated in mosquito antiviral defense^6–8^. The small interfering RNA (siRNA) pathway is considered the primary antiviral defense mechanism in dipteran insects^7,9–12^. The siRNA pathway relies on RNA interference (RNAi), an overarching mechanism of RNA silencing mediated by small RNAs (sRNAs). Viral double-stranded RNA (dsRNA) is recognized by Dicer2 (Dcr2), an endonuclease that cleaves viral RNA into siRNAs that are 21 nucleotides (nt) in length. Next, siRNA duplexes are loaded through a process that involves the dsRNA binding protein, R2D2^13^, into the Argonaute2 (Ago2)-containing RNA induced silencing complex (RISC), which subsequently targets viral RNA for cleavage. Dcr2 belongs to the family of the DExD/H-box helicases, similar to RIG-I-like receptors in vertebrates^14^.

The antiviral role of the siRNA pathway in *Ae. aegypti* has been established by several earlier studies. In the *Ae. aegypti* cell line Aag2, *Dcr2* knockout resulted in higher Semliki Forest virus replication, and *Dcr2* transient reexpression rescued the antiviral phenotype^15^. In adult *Ae. aegypti* mosquitoes, *Dcr2* knockdown increased Sindbis virus (SINV) and dengue virus (DENV) titers after an infectious blood meal^16,17^, and shortened the extrinsic incubation period (i.e., the time required for an infected mosquito to become infectious) before DENV transmission^17^. However, a few studies indicated that the antiviral activity of the siRNA pathway in *Ae. aegypti* could not always be generalized. Antiviral activity of *Dcr2* upon DENV infection was observed in different laboratory strains^17^ but not in a field-derived mosquito strain^18^. This could reflect strain-specific differences in the contribution of antiviral immune pathways to DENV infection or could be linked to technical variation such as differences in gene knockdown efficiency. Elevated levels of SINV replication were observed upon midgut-specific *Dcr2* knockdown, suggesting an antiviral role of the siRNA pathway in this tissue^19^. In contrast, another study showed that the siRNA pathway failed to limit DENV replication in the mosquito midgut, although it efficiently restricted systemic DENV replication and remained effective in the midgut against both endogenous and exogenous dsRNA substrates^20^. This could be linked to the dsRNA-binding protein Loqs2 (a paralog of Loquacious and R2D2 that is not expressed in the midgut), because transgenic expression of *Loqs2* in the midgut was sufficient to decrease DENV RNA levels upon an infectious blood meal^20^.

In addition to studies using gene-silencing assays to target the siRNA pathway, a single study to date reported an attempt to mutate *Dcr2* using transcription activator-like effector nucleases (TALENs) in *Ae. aegypti* ^21^. In a subsequent study, an increase in yellow fever virus replication was found in the *Dcr2* mutant line, but no further phenotypic characterization of the line was provided^22^. More recently, the impact of the siRNA pathway on arbovirus replication in *Ae. aegypti* was assessed using transgenic overexpression of *Dcr2* and *R2D2* in the midgut following an infectious blood meal. Replication of DENV, Zika virus (ZIKV), and chikungunya virus (CHIKV) was suppressed upon overexpression of both siRNA genes, consistent with a broad-spectrum antiviral activity of the siRNA pathway in *Ae. aegypti* ^23^.

Despite the well-established antiviral activity of the siRNA pathway *in vivo* in *Ae. aegypti*, the full antiviral spectrum of the siRNA pathway and its effective contribution to transmission of arboviruses from different families remain patchy. Most earlier studies on the antiviral activity of the siRNA pathway in *Ae. aegypti* focused on measuring viral replication in various tissues rather than directly assessing virus transmission potential. Arbovirus transmission is not only influenced by vector competence, defined as the intrinsic ability of mosquito to acquire and subsequently transmit a pathogen^24^, but also by other entomological parameters such as survival rate, which has not been assessed in previous studies on the siRNA pathway in *Ae. aegypti*. While exploiting the antiviral activity of the siRNA pathway as a novel arbovirus control strategy in the future is very promising, characterizing the full impact of manipulating key siRNA pathway components on arbovirus transmission is an essential prerequisite.

Here, we report on the generation and initial characterization of a viable and fertile *Dcr2* null mutant line of *Ae. aegypti*. We monitored the infection dynamics of four medically-relevant arboviruses, namely two flaviviruses: DENV^25,26^ and ZIKV^27^; and two alphaviruses: CHIKV^28^ and Mayaro virus (MAYV)^29,30^. The lack of *Dcr2* modestly enhanced early viral replication in the midgut and promoted early viral dissemination from the midgut to other tissues. However, the absence of *Dcr2* did not result in major changes in the overall level of vector competence. Additionally, the survival of *Dcr2* mutants was strongly reduced after infection with CHIKV (an alphavirus) but not with DENV (a flavivirus). Together, our results define a multifaceted role for *Dcr2* in arbovirus transmission by *Ae. aegypti* mosquitoes.

## Results

### Generation of an *Ae. aegypti Dcr2* null mutant line

To investigate the role of *Dcr2* and the siRNA pathway during arbovirus infection in *Ae. aegypti*, we generated a *Dcr2* null mutant line using CRISPR/Cas9-mediated gene editing^31^. The line was generated from a field-derived *Ae. aegypti* colony originating from Gabon^32^. In brief, we injected mosquito embryos with the Cas9 endonuclease coupled to single-guide RNAs (sgRNAs) targeting exon 5 (that had previously been targeted with TALENs^21^) and exon 7 of the *Dcr2* gene (*AAEL006794*) and obtained generation zero (G_0_) adult mosquitoes with potential gene edits. We screened the *Dcr2* locus of the G_0_ individuals for mutations and found, in one male mosquito, an insertion of 2 base pairs (bp) followed by point mutations of the next 2 bp in exon 5 (see Methods for details). The sequence edit led to a frameshift and a new open reading frame in which a stop codon was introduced at position 174 of the Dcr2 protein sequence (Figure 1A). Thus, we generated a mutant line encoding a truncated version of the Dcr2 protein with only 173 of the 1658 amino acids. The first 173 amino acids of Dcr2 correspond to the DEAD component of the DEAD-box helicase domain, which is responsible for unwinding dsRNA and initiating RNAi. Because the DEAD-box helicase domain was left incomplete, we hypothesized that this mutation triggered a *Dcr2* loss of function and called the mosquito mutant line *Dicer2^null^*. In parallel of generating the mutant line, we also produced a wild-type “sister” line derived from the same crossing scheme as the mutant line, which we refer to as the control line hereafter.

**Figure 1.**
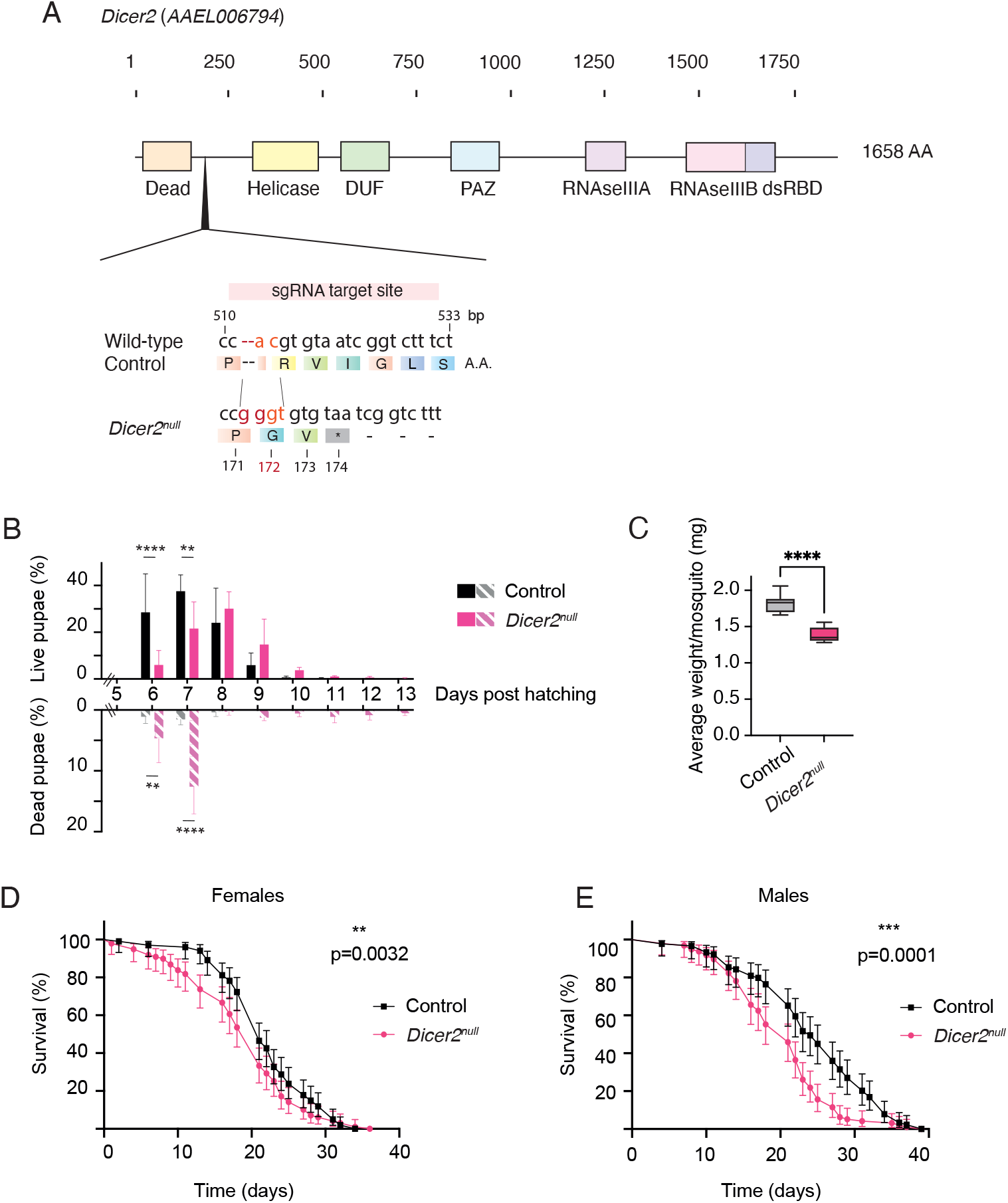
*Dicer2^null^* mutants display modest fitness defects. (**A**) Structure of the *Dicer2* locus (*AAEL006794*) with functional domains highlighted in colored boxes and size shown in amino-acid (AA) length. Nucleotide and AA sequence of the region targeted by the single-guide RNA are shown for *Dicer2^null^* mutants and wild-type mosquitoes. (**B**) Pupation success in the *Dicer2^null^* mutant and control lines. Numbers of live and dead pupae were scored daily and percentages were calculated relative to the total number of pupae present at the end of the experiment. Data are shown as mean and standard deviations (s.d.) of 3 independent groups of 150-200 pupae each. Statistical significance of the pairwise differences was determined with Student’s t-test (*p<0.05; **p<0.01; ***p<0.001). (**C**) Average weight of *Dicer2^null^* mutant and control adult female mosquitoes. Data are shown as means and s.d. of six replicates containing 5 mosquitoes that had been starved 24 hours before weighing. Statistical significance of the differences was determined with Student’s t-test (*p<0.05; **p<0.01; ***p<0.001). (**D**, **E**) Survival curves of female (**D**) and male (**E**) adult *Dicer2^null^* mutants and controls under standard insectary conditions. Mosquitoes were anesthetized on ice and sorted in incubation boxes by groups of 25 in 4 replicates for each condition. Mortality was scored daily and fresh pads containing 10% sucrose solution were renewed twice a weekly. Statistical significance of the differences was determined by Gehan-Breslow-Wilcoxon tests (*p<0.05; **p<0.01; ***p<0.001).

### *Dicer2^null^* mutants display moderate fitness defects

The final step in establishing the *Dicer2^null^* mutant and control lines consisted of crossing heterozygote individuals (carrying one copy of the mutant *Dcr2* allele and one copy of the wild-type allele) to obtain homozygous mutants (the mutant line) and homozygous wild-type individuals (the control “sister” line). Although the *Dicer2^null^* mutant line was viable and fertile, we detected a series of modest fitness defects relative to the control line. First, we measured daily pupal mortality over the 9-day time window during which pupation is expected to occur in our laboratory conditions. We found that pupal death rates were higher in *Dicer2^null^* mutants than in the control line on day 6 and 7 post hatching (Figure 1B). For instance, on day 7, we observed 12.5% pupal mortality in the *Dicer2^null^* line, compared to only 1.5% in the control line (p<0.0001). The cumulative pupal death rate was of 22.1% for *Dicer2^null^* mutants compared to 3.1% for controls. Finally, we used a four-parameter logistic model to determine the median pupation time which was of 7.5 days for *Dicer2^null^* mutants compared to 6.4 days for control mosquitoes (p=0.0002). Thus, we concluded that *Dicer2^null^* mutants had slower larval development and higher pupal mortality.

Second, we measured the weight of adult female mosquitoes of 5-7 days of age. We starved mosquitoes 24h prior to weighing to ensure that no sugar water remained in their digestive tract. We found that *Dicer2^null^* mutants had a lower body weight (1.3 mg/mosquito on average) than the control line (1.8 mg/mosquito; p<0.0001), despite exactly similar rearing conditions (Figure 1C). Altogether, these results point to fitness defects during development in the *Dicer2^null^* line.

Next, we tested whether adult lifespan was affected in the *Dicer2^null^* mutant line by performing a survival analysis in male and female adult mosquitoes kept on a sugar-only diet in standard rearing conditions. We found a slight reduction in survival for *Dicer2^null^* females (median lifespan of 21 days vs. 22 days in controls; p=0.02; Figure 1D) and stronger reduction in survival for *Dicer2^null^* males (median lifespan of 21 days vs. 24 days in controls; p<0.0001; Figure 1E). Altogether, our data demonstrate moderate fitness losses in developing and adult *Dicer2^null^* mutants.

### The siRNA pathway is impaired in *Dicer2^null^* mutants

To assess whether the siRNA pathway was impaired in *Dicer2^null^* mutant mosquitoes, we first used an *in vivo* RNAi reporter assay^33^. In brief, adult females were injected intrathoracically with a transfection mixture containing a Firefly luciferase reporter plasmid and dsRNA targeting Firefly luciferase (Fluc) or Green Fluorescent Protein (GFP) as a negative control. A reporter plasmid encoding a Renilla luciferase (Ren) was also added and used as an *in vivo* transfection control. Three days later, whole mosquito bodies were homogenized, and Firefly and Renilla luciferase luminescence counts were measured (Figure 2A). Results were expressed as log_10_-transformed Fluc/Ren ratios and we analyzed the data from two separate experiments by accounting for the variation between experiments (see Methods for details). When injected with reporter plasmids and dsRNA targeting GFP, *Dicer2^null^* mutant and control lines displayed comparable Fluc/Ren ratios, confirming that Firefly and Renilla luciferase were well translated, and that non-specific silencing activity was not detected (Figure 2B). However, when *Dicer2^null^* mutant and control lines were injected with reporter plasmids and dsRNA targeting Fluc, the luciferase counts were significantly higher in *Dicer2^null^* mutants (Figure 2B). This indicates that Fluc expression from the plasmid was not efficiently silenced, and that RNAi activity was in fact reduced in *Dicer2^null^* mutants.

**Figure 2.**
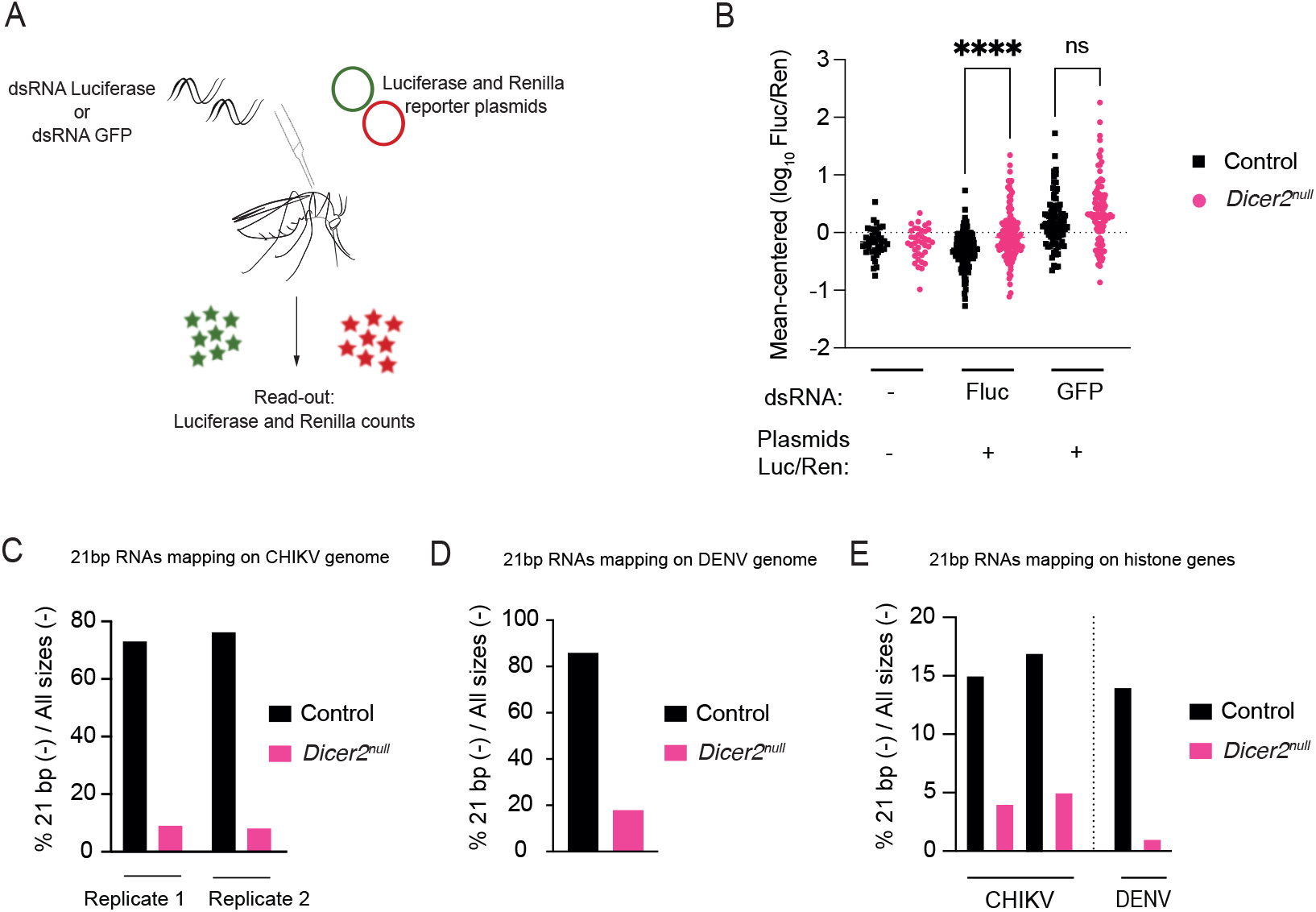
The siRNA pathway is impaired in *Dicer2^null^* mutants. (**A**) Experimental scheme of the RNAi reporter assay. Adult *Dicer2^null^* mutants and control mosquitoes were co-injected with Firefly luciferase (Fluc) and Renilla luciferase (Rluc) reporter plasmids, and dsRNA targeting Fluc or GFP. Luminescence was measured 3 days post injection in individual mosquitoes. (**B**) *In vivo* RNAi activity in the *Dicer2^null^* mutant and control lines. Luminescence counts of Firefly luciferase were normalized to the Renilla luciferase counts and log_10_ transformed. Mean-centered values represent the raw data corrected for differences between two experimental replicates for visual purposes. Mean-centered values were compared by one-way ANOVA (*p<0.05; **p<0.01; ***p<0.001). (**C, D, E**) Production of siRNAs in the *Dicer2^null^* mutant and control lines. Female *Dicer2^null^* mutants and control mosquitoes were exposed to CHIKV or DENV-1, and mosquito carcasses were harvested on day 5 (CHIKV, in duplicates) or day 6 (DENV-1) post infection. Virus infection was confirmed by RT-qPCR prior to sRNA sequencing. Negative 21-bp RNA reads were mapped to CHIKV (**C**), DENV-1 (**D**), or a histone transcript cluster located between positions 161,180,000 and 161,236,000 of chromosome 3 (**E**), and mapped reads expressed as a percentage relative to all sizes of negative reads.

Next, we aimed to assess siRNA production levels by performing high-throughput sRNA sequencing of *Dicer2^null^* mutant and control lines exposed to DENV-1 and CHIKV via an infectious blood meal. Five days post virus exposure, we extracted RNA from mosquito carcasses (i.e., abdomen without the midgut) and confirmed their infected status by RT-qPCR before preparing sRNA sequencing libraries. We found that indeed, the percentage of 21-bp RNAs mapping on the CHIKV genome (Figure 2C) or the DENV-1 genome (Figure 2D) was strongly reduced in *Dicer2^null^* mutants compared to controls. Additionally, we assessed the production of endogenous siRNAs in the same mosquitoes infected with DENV-1 or CHIKV. We found that the percentage of 21-bp RNAs mapping on a cluster of histone genes was strongly reduced in *Dicer2^null^* mutants (Figure 2E). Taken together, our results demonstrate that both exogenous and endogenous siRNAs are undetectable in *Dicer2^null^* mutants.

### *Dicer2* impacts viral infection dynamics upon alphavirus infection *in vivo*

We performed a series of experimental infections aimed at analyzing the impact of *Dcr2* on viral infection dynamics *in vivo*. We exposed *Dicer2^null^* mutant and control lines to blood meals containing alphaviruses (CHIKV and MAYV; Figure 3) or flaviviruses (DENV and ZIKV; Figure 4), and dissected the midgut, carcass and head at two time points post exposure chosen to represent early and late stages of the infection process. At the late time point, we also collected mosquito saliva to assess the impact of *Dcr2* on virus transmission potential, a direct measure of vector competence. For every time point and body part, we assessed infection prevalence and viral loads, by RT-qPCR (DENV) or infectious titration assay (CHIKV, MAYV and ZIKV). Midgut prevalence was calculated as the number of virus-positive midguts over the total number of exposed mosquitoes. Carcass prevalence was calculated as the number of virus-positive carcasses over the number of virus-positive midguts. Head prevalence was calculated as the number of virus-positive heads over the number of virus-positive carcasses. Finally, saliva prevalence was calculated as the number of virus-positive saliva samples over the number of virus-positive heads. To quantify overall vector competence, we also calculated the proportion of virus-exposed mosquitoes with virus-positive saliva, which we refer to as transmission efficiency (Figure 5). When saliva testing could not be performed, we used the proportion of virus-exposed mosquitoes with a virus-positive head as a proxy for transmission efficiency.

**Figure 3.**
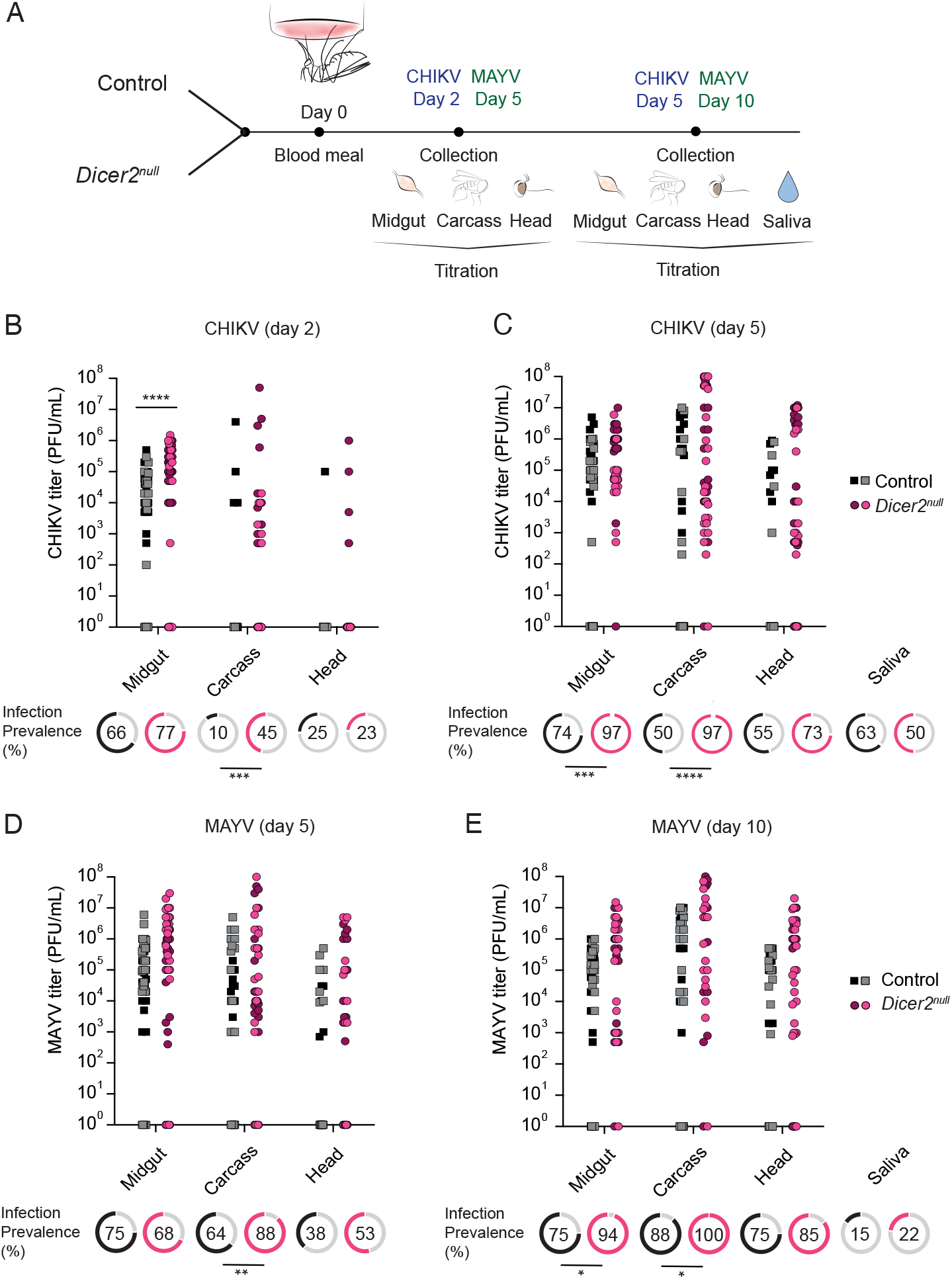
*Dcr2* hinders early infection dynamics of alphaviruses. (**A**) Experimental scheme for the analysis of CHIKV and MAYV infection dynamics. Adult *Dicer2^null^* mutants and control female mosquitoes were exposed to an infectious blood meal containing 10^6^ PFU/mL of CHIKV or MAYV. Midguts, carcasses, and heads from the same mosquito were collected on day 2 and 5 post exposure (CHIKV) or day 5 and 10 post exposure (MAYV). Saliva was only collected 5 (CHIKV) or 10 (MAYV) days post exposure. Infection prevalence and CHIKV titers were determined by plaque assay 2 (**B**) and 5 (**C**) days post exposure. Infection prevalence and MAYV titers were determined by plaque assay 5 (**D**) and 10 (**E**) days post exposure. The viral loads from two experimental replicates are combined and depicted using two shades of the same color (grey/black for control mosquitoes, pink/red for *Dicer2^null^* mutants). The data are merged for infection prevalence because the experiment effect was non-significant. Each experimental condition is represented by 48-56 mosquitoes (**B**, **C**; CHIKV) or 37-64 mosquitoes (**D**, **E**; MAYV). The statistical significance of pairwise comparisons shown in the figure (*p<0.05; **p<0.01; ***p<0.001) was obtained with a Student’s t-test applied to the raw values after correction for the variation between experiments (viral loads) or a Chi^2^ test (prevalence). The full statistical analyses are provided in Supplementary Figure 1A (CHIKV) and Supplementary Figure 1B (MAYV).

**Figure 4.**
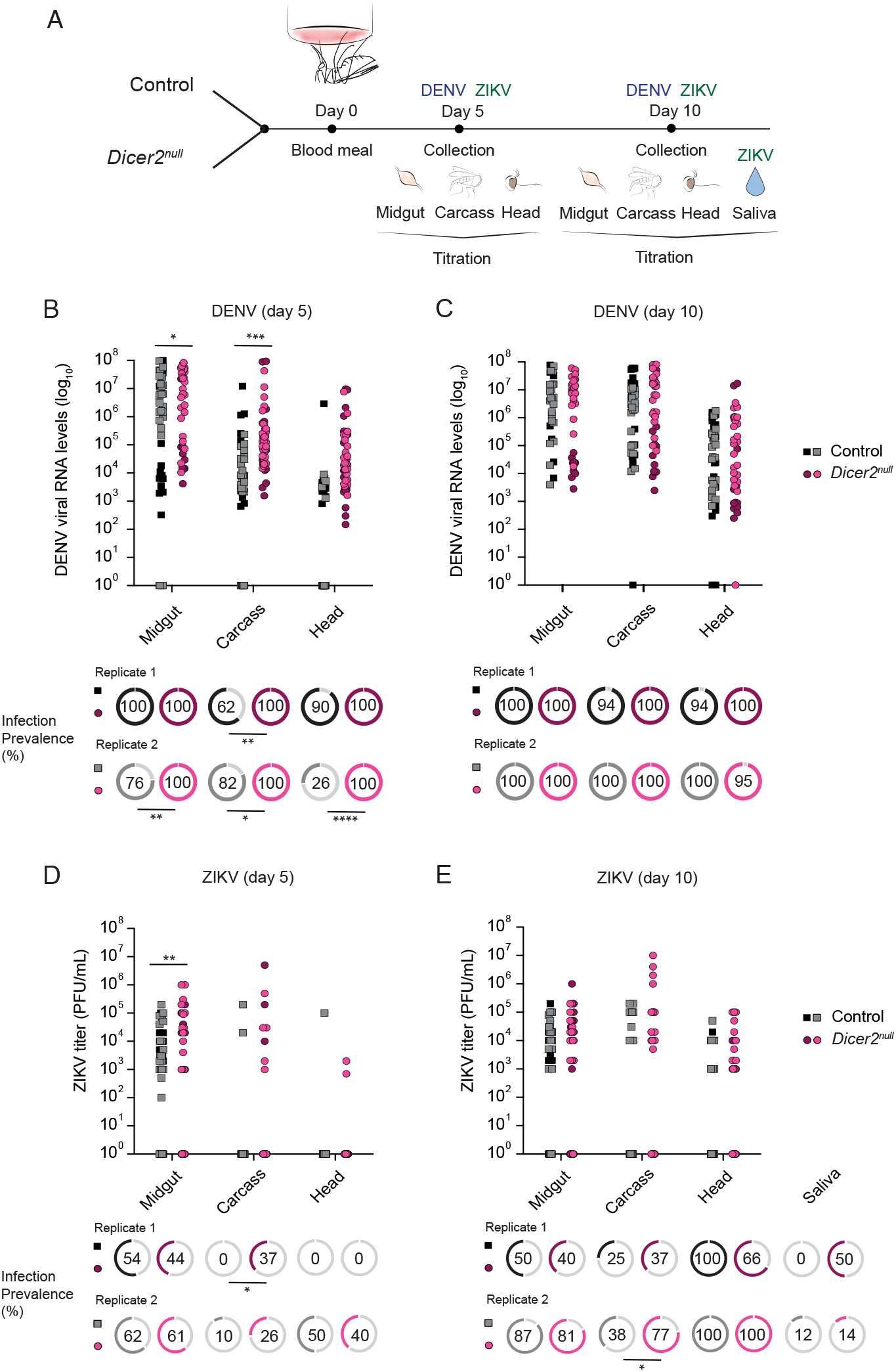
*Dcr2* hinders early infection dynamics of flaviviruses. (**A**) Experimental scheme for the analysis of DENV-3 and ZIKV infection dynamics. Adult *Dicer2^null^* mutants and control female mosquitoes were exposed to an infectious blood meal containing 5.10^6^ FFA/mL of DENV-3 or 10^6^ PFU/mL of ZIKV. Midguts, carcasses and heads from the same mosquito were collected on day 5 and 10 post exposure. Saliva was only collected 10 days post exposure with ZIKV. Infection prevalence and DENV-3 titers were determined by RT-qPCR 5 (**B**) and 10 (**C**) days post exposure. Infection prevalence and ZIKV titers were determined by plaque assay 5 (**D**) and 10 (**E**) days post exposure. Data from two experimental replicates are combined and depicted on the graph using two shades of the same color (grey/black for control mosquitoes, pink/red for *Dicer2^null^* mutants). Each experimental condition is represented by 40-50 mosquitoes (**B**, **C**; DENV-3) or 42-56 mosquitoes (**D**, **E**; ZIKV). The statistical significance of pairwise comparisons shown in the figure (*p<0.05; **p<0.01; ***p<0.001) was obtained with a Student’s t-test after correction for the variation between experiments (viral loads) or a Chi^2^ test (prevalence). The full statistical analyses are provided in Supplementary Figure 2A (DENV-3) and Supplementary Figure 2B (ZIKV).

**Figure 5.**
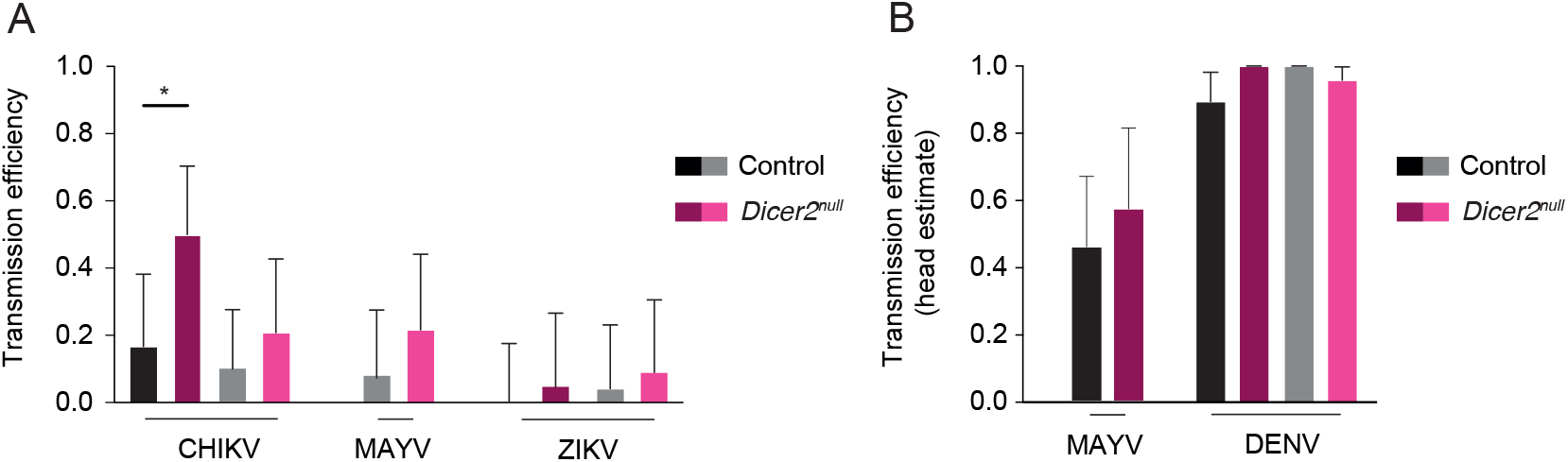
*Dcr2* does not significantly impact overall vector competence for arboviruses. (**A**) Transmission efficiency (the overall proportion of virus-exposed females with virus-positive saliva) is shown for each experimental replicate of CHIKV (Figure 3), ZIKV (Figure 4) and the second replicate of MAYV infection (Figure 3). (**B**) A proxy for transmission efficiency (the overall proportion of virus-exposed females with virus-positive heads) is shown for both experimental replicates of DENV-3 infection (Figure 4) and the first replicate of MAYV infection (Figure 3) because saliva samples were not available. Statistical significance of the difference in proportions was determined with Fisher’s exact test (*p<0.05; **p<0.01; ***p<0.001).

First, we exposed *Dicer2^null^* mutants and control mosquitoes to an infectious blood meal containing 10^6^ plaque-forming units (PFU)/mL of CHIKV, and collected body parts (midgut, body, head) 2 and 5 days post exposure, as well as saliva at the later time point (Figure 3A). Two days post exposure, we found a significantly higher midgut viral titer in *Dicer2^null^* mutants, but no difference in infection prevalence (Figure 3B). We found a significantly higher carcass prevalence in *Dicer2^null^* mutants compared to controls (45% vs. 10%; p=0.0008), but no difference in infectious titers in the virus-positive carcasses. Five days after the infectious blood meal, we did not detect any differences in viral loads in any body part (Figure 3C). However, we observed a higher midgut prevalence (97% vs. 74%; p=0.0007) and higher carcass prevalence (97% vs. 50%; p<0.0001) in *Dicer2^null^* mutants relative to controls. Infection prevalence in heads and saliva samples did not significant differ between *Dicer2^null^* mutants and controls on day 5 post exposure. To determine the overall impact of *Dcr2* on vector competence, we compared the transmission efficiency of the *Dicer2^null^* mutant and control lines on day 5 (Figure 5A). *Dicer2^null^* mutants had a marginally significantly higher transmission efficiency in the first experiment (50% vs. 17%; p=0.0305) but not in the second experiment (21% vs. 10%; p=0.4429). These results demonstrate that the loss of *Dcr2* enhances early CHIKV replication in the midgut, promotes early viral dissemination from the midgut to the abdomen, and results in a higher midgut infection rate and higher dissemination rate from midguts to abdomens on day 5 post exposure. However, these differences do not lead to a major increase in vector competence on day 5 post virus exposure.

To test whether such observations could be extended to another virus of the *Alphavirus* genus, we exposed mosquitoes to an infectious blood meal containing 10^6^ PFU/mL of MAYV and collected samples 5 and 10 days post exposure (Figure 3A). On day 5, the viral loads in midguts, carcasses and heads did not differ between *Dicer2^null^* and control mosquitoes (Figure 3D). Infection prevalence was not altered in the mosquito midguts or heads, but it increased significantly in carcasses of *Dicer2^null^* mutants relative to controls (88% vs. 64%; p=0.0069). On day 10 post exposure, we did not detect differences in viral loads between mosquito lines in any body part but observed a moderate increase of infection prevalence in midguts (from 75% to 94%; p=0.0157) and carcasses (from 88% to 100%; p=0.0424) in *Dicer2^null^* mutants relative to controls (Figure 3E). We found no difference in infection prevalence in heads and saliva samples on day 10 post exposure. Our estimates of transmission efficiency were based on virus-positive heads in the first experiment (Figure 5B) and virus-positive saliva samples in the second experiment (Figure 5A). Transmission efficiency estimates on day 10 did not significantly differ between *Dicer2^null^* mutants and controls in the first experiment (57% vs. 45%; p=0.7374) or in the second experiment (22% vs. 8%; p=0.2448). These results demonstrate that *Dcr2* does not impact MAYV viral loads 5 and 10 days post exposure in the mosquito midgut, carcass, or head. Yet, loss of *Dcr2* increased MAYV carcass prevalence on day 5 and 10, and midgut prevalence on day 10, similar to our observation with CHIKV. Thus, we conclude that *Dcr2* does not seem to play a major role in the replication of the alphaviruses we tested but contributes to controlling viral dissemination from the midgut to the carcass. Overall, these differences do not result in a major change in vector competence on day 10 post virus exposure.

### *Dicer2* impacts viral infection dynamics upon flavivirus infection *in vivo*

To evaluate the impact of *Dcr2* on flavivirus infection dynamics, we next exposed mosquitoes to an infectious blood meal containing 5 × 10^6^ focus-forming units (FFU)/mL of DENV-3 and collected samples 5 and 10 days post exposure (Figure 4A). Overall, infection prevalence was higher in the first experimental replicate and we accounted for this effect in the statistical model and by plotting infection prevalence separately. Five days after exposure, infection prevalence was significantly higher in *Dicer2^null^* mutants for all body parts (midgut, carcass and head) in the second experimental replicate. In the first replicate, the difference in infection prevalence was only significantly higher in the carcass of *Dicer2^null^* mutants (100% vs. 62%; p=0.0052), but our ability to detect differences was limited by the higher overall infection prevalence in this experiment (Figure 4B). We concluded that DENV-3 dissemination from the midgut to the abdomen, and from the abdomen to the head was enhanced in *Dicer2^null^* mutants. This was associated with an increase of viral RNA loads in the midguts (p=0.0230) and carcasses (p=0.0006) of *Dicer2^null^* mutants on day 5 post exposure (Figure 4B). Ten days post exposure, infection prevalence was almost 100% in both experimental replicates and all body parts, indicating that DENV-3 disseminated throughout the mosquito body (Figure 4C). We did not observe any differences in viral RNA levels between *Dicer2^null^* mutants and control mosquitoes on day 10 in any body part, when viral levels are high and most likely reaching a plateau. To compare transmission efficiency between the *Dicer2^null^* mutant and control lines, we used the proportion of virus-exposed mosquitoes with a virus-positive head because saliva results were unavailable (Figure 5B). *Dicer2^null^* mutants did not exhibit significantly higher transmission efficiency on day 10 in the first experiment (100% vs. 89%; p=0.4891) or in the second experiment (96% vs. 100%; p=0.9999). Overall, our analysis of DENV-3 infection dynamics demonstrates that *Dcr2* restricts early viral dissemination from the midgut to the abdomen, and from the abdomen to the head, probably by controlling early viral replication in the midgut and carcass.

Finally, we proceeded to analyze the impact of *Dcr2* on infection dynamics of another member of the *Flavivirus* genus, ZIKV. We collected samples on day 5 and 10 after exposure to an infectious blood meal containing 10^6^ PFU/mL of ZIKV (Figure 4A). Overall infection prevalence differed between the two experimental replicates, and we accounted for this effect in the statistical model and by displaying the data separately. We only found subtle differences in ZIKV infection dynamics between *Dicer2^null^* mutants and control mosquitoes. Infectious titers were higher in the midguts of *Dicer2^null^* mutants on day 5 (Figure 4D). Carcass prevalence was higher in *Dicer2^null^* mutants than in control mosquitoes only on day 5 in the first experimental replicate (37% vs. 0%; p=0.0171; Figure 4D) and on day 10 in the second experimental replicate (77% vs. 38%; p=0.0127; Figure 4E). We determined the overall impact of *Dcr2* on vector competence by estimating transmission efficiency on day 10 (Figure 5A). Transmission efficiency did not significantly differ between the *Dicer2^null^* mutant and control lines in the first experiment (5% vs. 0%; p=0.4545) or in the second experiment (9% vs. 4%; p=0.6). We concluded that the loss of *Dcr2* enhanced early ZIKV replication in the midgut, and facilitated viral dissemination to the abdomen, but this did not result in a detectable change in vector competence on day 10 post virus exposure.

Taken together, our data on the infection dynamics of CHIKV, MAYV, DENV, and ZIKV reveal a role for *Dcr2* in controlling early viral replication and dissemination, when viruses are infecting the midgut and starting to disseminate systemically in the abdomen. We found that the loss of *Dcr2* increased viral loads in the midgut (except for MAYV) and promoted viral dissemination from the midgut to the carcass (except for ZIKV) at early infection time points. Conversely, our data showed that *Dcr2* was not required for controlling viral replication at later infection stages, at least in the body parts that we analyzed. The overall level of vector competence for the four arboviruses was similar between the *Dicer2^null^* mutant line and the control line (Figure 5), except for one CHIKV experiment.

### *Dicer2* impacts viral replication of CFAV *in vivo*

In addition to arboviruses, we asked whether *Dcr2* could impact replication levels of cell-fusing agent virus (CFAV), an insect-specific flavivirus^34^. We inoculated mosquitoes intrathoracically with 50 tissue-culture infectious dose 50% (TCID_50_) units of CFAV and measured viral RNA levels by RT-qPCR in whole mosquito bodies 2, 4, 6 and 8 days post infection (Supplementary Figure 3A). We found that *Dcr2* had a very limited impact on CFAV loads. We only observed slightly increased viral loads on days 2 and 6 post inoculation in *Dicer2^null^* mutants compared to control mosquitoes (Supplementary Figure 3B), indicating a limited influence of *Dcr2* on the systemic replication of the insect-specific flavivirus CFAV. This is consistent with our observations above demonstrating a limited role for *Dcr2* in controlling systemic replication of arboviruses.

### *Dicer2* prevents virus-induced mortality upon CHIKV, but not DENV infection

Finally, we asked whether *Dcr2* could impact mosquito survival after exposure to arboviruses via an infectious blood meal. We monitored the survival of *Dicer2^null^* mutants and control mosquitoes after oral exposure to 10^6^ PFU/mL of CHIKV or 5 × 10^6^ FFU/mL of DENV-3 and included a non-infectious blood meal as a negative control. We verified on day 7 post blood meal that most of the mosquitoes (>94% across conditions) were virus-positive (Supplementary Figure 4). The survival rates of control mosquitoes did not differ after CHIKV exposure or mock exposure (p=0.09; Figure 6A). In contrast, the survival rate of *Dicer2^null^* mutants was significantly reduced after CHIKV exposure relative to the mock exposure control (p<0.0001; Figure 6B). Median survival time was 10 days upon CHIKV infection, compared to 18.5 days upon mock infection. Thus, we concluded that *Dcr2* was required to prevent virus-induced mortality during CHIKV infection. For DENV-3, the survival rates of control mosquitoes slightly increased after virus exposure relative to mock exposure (p=0.01; Figure 6C). However, we did not detect any difference in the survival rates of *Dicer2^null^* mutants after DENV-3 exposure or mock infection (p=0.63; Figure 6D). These data indicate that *Dcr2* does not influence mosquito survival during infection with DENV-3. Taken together, our survival analyses reveal different consequences of *Dcr2* absence on mosquito survival upon DENV-3 or CHIKV infection. Whereas *Dcr2* is required to prevent virus-induced mortality upon CHIKV infection, this is not the case upon DENV-3 infection.

**Figure 6.**
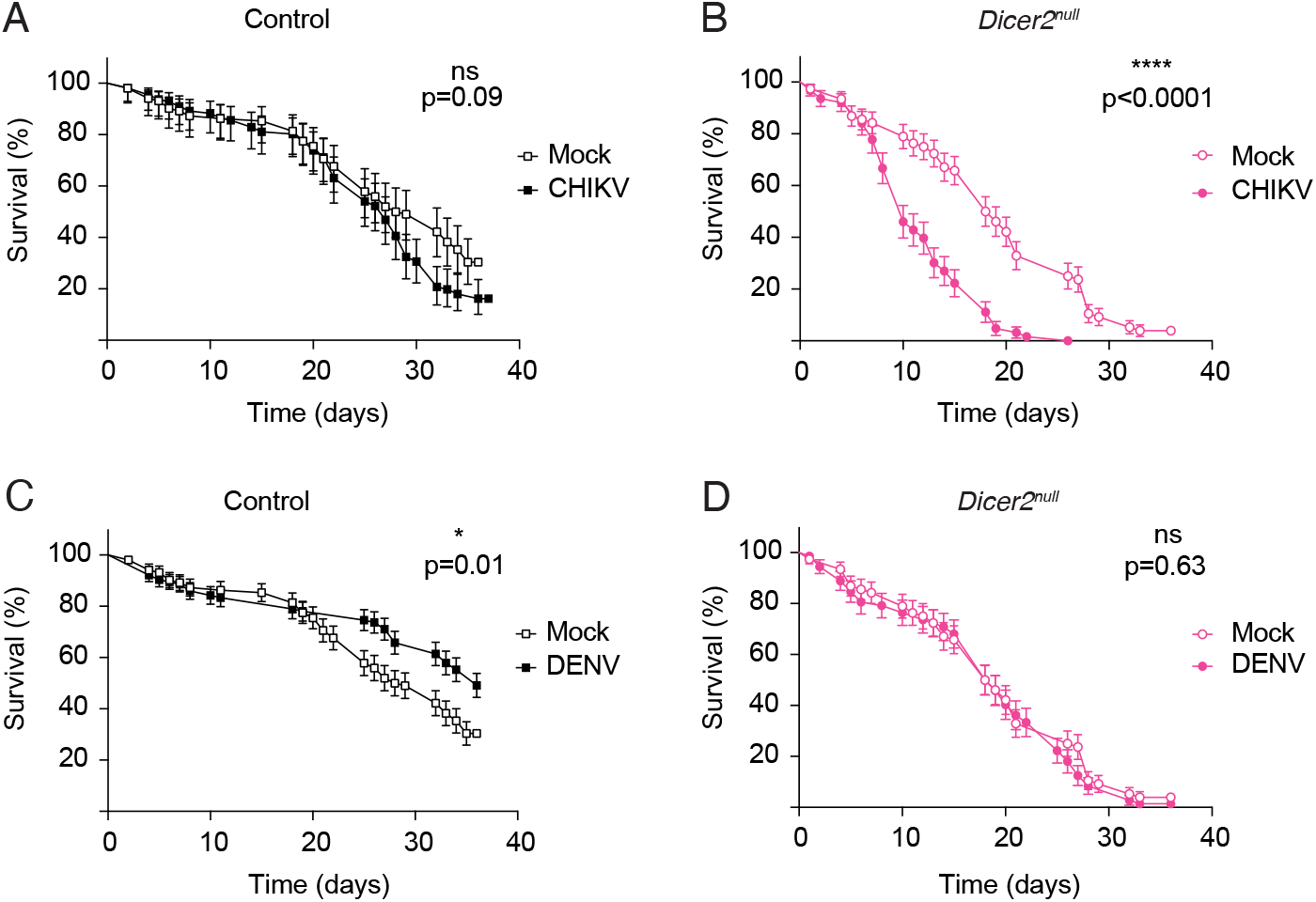
*Dcr2* prevents virus-induced mortality during CHIKV infection but not DENV-3 infection. Adult female mosquitoes of the control line (**A**, **C**) or *Dicer2^null^* mutant line (**B**, **D**) were exposed to an infectious blood meal containing 10^6^ PFU/mL of CHIKV (**A**, **B**), 5 × 10^6^ FFU/mL of DENV-3 (**C**, **D**), or a non-infectious blood meal (mock). Data are shown as mean and s.d. of 3 replicates represented by 21-38 mosquitoes each. Mortality was scored daily and fresh pads containing 10% sucrose solution were renewed bi-weekly. Statistical significance of the differences was determined by Kaplan-Meier analysis and Gehan-Breslow-Wilcoxon tests (*p<0.05; **p<0.01; ***p<0.001). See also Supplementary Figure 4 for corresponding mosquito infection levels.

## Discussion

The siRNA pathway is considered a major, broad-spectrum antiviral pathway in insects^35^. The antiviral activity of the siRNA pathway is well established in mosquito vectors such as *Ae. aegypti*, and thus represents a potential key target in the development of novel arbovirus control strategies aimed at developing lab-engineered mosquitoes uncapable of transmitting viral pathogens. Several earlier studies have characterized the molecular components and function of the siRNA pathway in *Ae. aegypti*, but surprisingly little is known on its effective impact on arbovirus transmission. Here, we assessed the effect of genetically inactivating the siRNA pathway on infection dynamics and transmission potential of several arboviruses of medical significance.

Our *Dicer2^null^* mutant line is viable and fertile, demonstrating that *Dcr2* is not an essential gene in *Ae. aegypti*, at least not in the genetic background of our Gabonese mosquito colony. The *Dicer2^null^* mutant line does exhibit modest fitness defects such as slower development, increased pupal mortality and smaller adult body size. How exactly *Dcr2* relates to these fitness defects is currently unknown and deserves further investigation. We confirmed by sRNA sequencing that the *Dicer2^null^* mutant line does not produce siRNAs and that dsRNA-triggered silencing activity is in fact abolished. Surprisingly, we found that the loss of *Dcr2* had a limited impact on viral replication and infection dynamics of arboviruses acquired via an infectious blood meal. In fact, the antiviral effect of *Dcr2* could only be detected at the early stage of midgut infection and dissemination out of the midgut. *Dcr2* was not required for controlling viral replication during the later stage of infection when the virus has already disseminated throughout the mosquito body. Accordingly, the absence of *Dcr2* did not significantly affect systemic replication of the insect-specific virus CFAV following intrathoracic inoculation. We did not detect major changes in vector competence, estimated as the overall proportion of virus-exposed mosquitoes that eventually became infectious (transmission efficiency). Note that some of our transmission efficiency estimates could have been overestimated because they relied on virus detection in head tissues, not in saliva, however our comparisons between the mutant and control lines were always based on the same measurement. Additionally, the loss of *Dcr2* significantly increased virus-induced mortality during CHIKV infection, but not DENV-3 infection. Reduced survival of infected mosquitoes is predicted to strongly decrease arbovirus transmission, because only long-lived mosquitoes can effectively contribute to it^24^.

Our study demonstrated a modest but detectable impact of *Dcr2* on the control of arbovirus replication *in vivo* in *Ae. aegypti* mosquitoes. Most previously published studies relied on the use of gene knockdown assays to assess the contribution of siRNA pathway components to virus replication control^16–18^. The gene knockdown approach is limited by its intrinsic variability and efficiency in targeting gene expression. Thus, some of the previous studies were inconclusive because they could not disambiguate insufficient knockdown efficiency from a true lack of biological effect. Our *Dicer2^null^* mutant line allowed us to rule out such technical limitations and assess the effective contribution of *Dcr2* to the control of viral replication. Our results conclusively show that the impact of *Dcr2* on viral replication is transient, and mainly limited to the midgut tissue. A previous study reported that the siRNA pathway was not required for controlling DENV replication in the midgut^20^, based on gene knockdown of *Ago2*. It is possible that gene knockdown did not provide sufficient sensitivity to detect the minor contribution of the siRNA pathway to control DENV replication in the midgut. Alternatively, it is possible that the antiviral role we observed for *Dcr2* in the midgut is independent of the downstream factors of the siRNA pathway. Dcr2 is an RNA sensor that acts upstream of Ago2 and is possibly involved in other branches of antiviral immunity that impact viral replication in the midgut^18,23,36–38^.

Despite its modest impact on viral titers, we found that *Dcr2* had a significant role in controlling the rate of viral dissemination from the midgut to the rest of the mosquito body. We found that the absence of *Dcr2* promoted viral spread from the midgut to the mosquito hemocoel, as indicated by a higher proportion of virus-positive carcasses during the early stage of infection. This is consistent with previous studies reporting that silencing *Dcr2* and *Ago2* accelerated systemic dissemination of DENV^17,20^. The mechanistic basis of this observation is still unclear, but we speculate that for some viruses, such as CHIKV, the slight increase in midgut viral titers we observed in the *Dicer2^null^* mutant line may allow an earlier viral exit of the midgut tissue. However, higher rates of early dissemination were also observed in the case of MAYV infection, in the absence of increased midgut viral titers. The mechanisms of virus spread from the midgut to the hemocoel remain unknown and might involve active virus replication in circulating cell populations such as hemocytes^39^. The putative role of *Dcr2* in controlling virus replication in these cells, in particular phagocytes, remains to be investigated.

To our knowledge, our study is the first to describe a protective role for *Dcr2* against virus-induced mortality during arbovirus infection of *Ae. aegypti*. Virus-induced mortality is generally modest during arbovirus infection of mosquitoes^40,41^ and indeed we found that CHIKV or DENV infection had no major effect on the survival curves of control mosquitoes. In the absence of *Dcr2*, however, the median lifespan of mosquitoes was almost halved following CHIKV infection, but not DENV infection. Whether this discrepancy is virus strain-specific or genus-specific remains to be determined. In the case of CHIKV, the increased viral loads observed in the midgut of mutant mosquitoes during the early stage of infection could be directly responsible for the higher virus-induced mortality, due to increased tissue damage, for example. Alternatively, *Dcr2* could be necessary to prevent virus-induced mortality without directly acting on viral replication, a defense mechanism often referred to as tolerance^42^. This could be through activating signaling pathways that enhance tissue repair^43^ or limiting tissue damage upon infection, such as apoptosis^44^. The contrasted outcome observed with DENV could reflect differences in the physiopathology of DENV infection in mosquito tissues, or differences in the immune or tissue repair pathways activated by viral infection.

It is worth noting that in the case of CHIKV, the protective effect of *Dcr2* against virus-induced mortality benefits both the host and the virus. Indeed, viral fitness is positively correlated with mosquito lifespan because the amount of transmission depends on the probability that (1) the mosquito survives through the extrinsic incubation period, and (2) the life expectancy of the mosquito after becoming infectious^45^. From this perspective, *Dcr2* could be considered as a “proviral” factor. Further studies are necessary to quantify the net effect of *Dcr2* on cumulative virus transmission, resulting from the combination of antiviral (delayed dissemination) and proviral (longer lifespan) effects.

Beyond the scope of the current study was assessing the implication of *Dcr2* in other sRNA pathways, such as the PIWI-interacting RNA (piRNA) pathway, which primarily functions to silence transposons in the germline. Recent studies revealed the existence of virus-derived piRNAs during infection of *Ae. aegypti* and *Ae. albopictus* mosquitoes by RNA viruses^11^. Crosstalk between the siRNA and piRNA pathways could occur and contribute to maintaining host fitness during viral infection, potentially through the integration of viral DNA forms of RNA viruses as endogenous viral elements, which are a source of piRNAs^35^. piRNA production occurs at late infection stages in *Ae. aegypti* upon DENV infection (day 14) or CHIKV infection (day 9)^20,46^. The present study analyzed sRNA populations only on day 5 and 6 post infection and did not allow detection of piRNAs, hence preventing us from studying the potential impact of *Dcr2* on piRNA biogenesis.

In conclusion, our study sheds new light on the effective contribution of *Dcr2* to arbovirus transmission by *Ae. aegypti* mosquitoes. Using a panel of arboviruses, we demonstrated the role of *Dcr2* in controlling early viral replication and in restricting early systemic viral spread for both flaviviruses and alphaviruses. Overall, however, *Dcr2* was not a major determinant of vector competence based on our estimates of transmission efficiency. We also discovered that *Dcr2* was involved in preventing virus-induced mortality during CHIKV infection, challenging the idea that *Dcr2* is exclusively an antiviral factor. Finally, we anticipate that our *Dicer2^null^* mutant line will be instrumental to advance basic research on the role of sRNA pathways in mosquito biology. It will be an important tool to investigate the mechanisms underlying essential processes such as transposon control, antiviral defense in the germline, and possibly immune priming and memory.

## Data availability

The sRNA sequencing data were deposited to the Sequence Read Archive (accession number pending).

## Acknowledgements

We thank the Lambrechts lab and Saleh lab members for helpful comments and discussions. We are grateful to Catherine Lallemand for assistance with mosquito rearing, and to Catherine Bourgouin for access to the Femtoject injector. We thank Davy Jiolle and Christophe Paupy for providing the DENV-3 strain and the World Reference Center for Emerging Viruses and Arboviruses at the University of Texas Medical Branch for providing the ZIKV strain. We thank Diego Ayala, Davy Jiolle and Christophe Paupy for the initial field sampling of mosquitoes in Bakoumba, Gabon. We thank Alain Kohl and Emilie Pondeville for sharing the plasmids for the reporter assay.

## Authors contributions

S.H.M., A.B.C., A.H-L., V.G., A.B., H.B., L.F., M.-C.S. and L.L. designed the experiments; S.H.M., A.B.C., A.H-L., V.G., A.B. and H.B. performed the experiments; S.H.M., A.B.C., A.H.L., V.G., A.B., H.B., L.F., L.L. and M.-C.S. analyzed the data; S.H.M. and L.L. wrote the paper with input from co-authors. All authors reviewed and approved the final version of the manuscript.

## Methods

### Ethics statement

Mosquito artificial infectious blood meals were prepared with human blood. Blood samples were supplied by healthy adult volunteers at the ICAReB biobanking platform (BB-0033-00062/ICAReB platform/Institut Pasteur, Paris/BBMRI AO203/[BIORESOURCE]) of the Institut Pasteur in the CoSImmGen and Diagmicoll protocols, which had been approved by the French Ethical Committee Ile-de-France I. The Diagmicoll protocol was declared to the French Research Ministry under reference 343 DC 2008-68 COL 1. All adult subjects provided written informed consent. Genetic modification of *Ae. aegypti* was performed under authorization number 7614 from the French Ministry of Higher Education, Research and Innovation.

### Cells

C6/36 cells (derived from *Aedes albopictus*) were cultured in Leibovitz’s L-15 medium (Life Technologies) supplemented with 10% fetal bovine serum (FBS, Life Technologies), 1% non-essential amino acids (Life Technologies), 2% Tryptose Phosphate Broth (Life Technologies) and 1% Penicillin/Streptomycin (P/S, Life Technologies) at 28°C without additional CO_2_. Vero-E6 (ATCC CRL-1586) cells were cultured in Dulbecco’s modified Eagle’s medium (DMEM) GlutaMAX^®^ (Life Technologies) containing 10% FBS (Invitrogen) and 1% P/S (Life Technologies) at 37°C with 5% CO_2_.

### Virus strains

DENV-1 strain KDH0026A was originally isolated in 2010 from the serum of a patient in Kamphaeng Phet, Thailand^47^. DENV-3 strain GA28-7 was originally derived in 2010 from the serum of a patient in Moanda, Gabon^48^. ZIKV strain FSS13025 was originally isolated in 2010 from the serum of a patient in Cambodia^49^. CHIKV isolate M105 was originally derived in 2014 from the serum of a patient in Martinique^50^. MAYV strain TRVL 4675 was derived from a previously described infectious clone^51^. Viral stocks were prepared in C6/36 cells for DENV and ZIKV and Vero E6 cells for CHIKV and MAYV. The wild-type CFAV strain CFAV-KPP was isolated from mosquito homogenates and produced from an RNA template diluted in Opti-MEM™ Reduced Serum Medium (Gibco Thermo Fisher Scientific) as described^52^.

### Mosquito rearing

Mosquitoes were reared under controlled conditions (28°C, 12-hour light/12-hour dark cycle and 70% relative humidity) in an insectary. Prior to performing the experiments, their eggs were hatched synchronously in a SpeedVac vacuum device (ThermoFisher) for one hour. Their larvae were reared in plastic trays containing 1.5 L of tap water and supplemented with Tetramin (Tetra) fish food at a density of 200 larvae per tray. After emergence, adults were kept in BugDorm-1 insect cages (BugDorm) with permanent access to 10% sucrose solution.

### Mosquito genetic modification

The *Dicer2* gene (*AAEL006794*) was targeted using CRISPR/Cas9-mediated gene editing, using a protocol adapted from a previously described method^31^. Gene-targeting reagents were injected into a wild-type colony of *Ae. aegypti* mosquitoes originally from Bakoumba, Gabon^*32*^. Single-guide RNAs (sgRNAs) were designed using CRISPOR^53^ by searching for sgRNAs of 20 bp in length with the NGG protospacer-adjacent-motif (PAM). To reduce chances of off-target mutations, and give preference to highly specific sgRNAs, only guides with off-targets that contained 3 or more mismatches were selected. sgRNAs were produced exactly as previously described^31^. The final injection mix contained 300 ng/μL spCas9 protein (M0386, New England Biolabs), 40 ng/μL of each sgRNA (3 targeting exon 5 and 7 of *Dicer2* and 4 targeting the *kmo* gene, *AAEL008879*; sequences provided in Supplementary Table 1). Embryos were injected 30-60 min after egg laying using a FemtoJet (Eppendorf) equipped with microinjection quartz needles (QF100-70-10, Sutter Instrument), and handled according to a standard protocol^54^. Embryos were hatched 3 days post injection and reared to adulthood according to standard rearing procedures. When disrupted, *kmo* produces a mosaic red-eye phenotype in G_0_ adults. We used this phenotype to easily verify CRISPR/Cas9 activity and screen for *Dicer2* mutations in red-eyes G_0_ mosquitoes preferentially. Genomic DNA was extracted from single legs of individual mosquitos using NucleoSpin DNA Insect Kits (Machery-Nagel) or DNAzol DIRECT (Molecular Research Center, Inc.), according to manufacturer’s instructions. *Dicer2* mutations were first tracked by PCR using DreamTaq DNA Polymerase (Thermo-Fisher Scientific) and subsequent restriction enzyme assay of exon 5 (with BSaaI, NEB) and exon 7 (BSErI, NEB). PCR amplicons were also purified with MinElute PCR Purification Kit (Qiagen) and subjected to Sanger sequencing for sequence verification (sequences provided in Supplementary Table 1). The initial *Dicer2* mutant mosquito (G_0_) that founded the *Dicer2^null^* mutant line was a male individual with mosaic red eyes (indicating mutation in the *kmo* gene), and mutations in both exon 5 and 7 of the *Dicer2* gene. Only the mutation in exon 5 that led to an early stop codon was tracked throughout subsequent crosses. The G_0_ founder mutant male was outcrossed to 8 wild-type females from the Bakoumba colony. The first generation (G_1_) inherited the *Dicer2* mutation, but did not display the red-eye phenotype, indicating non-inheritance of the *kmo* mutation. A single G_1_ male heterozygous for the *Dicer2* mutation was outcrossed to 10 wild-type females. For the subsequent cross, a brother-sister mating was performed using 13 and 10 heterozygous G_2_ males and females, respectively. This cross was intended to be the final cross to establish two G_3_ lines: (1) a “sister line” composed of individuals homozygous for the wild-type allele to be used as a control for the crossing scheme, and (2) a homozygous *Dicer2* mutant line. At this point, the “sister line” was successfully established with 26 males and 43 females homozygous for the wild-type allele in exon 5 of *Dicer2* and maintained separately. Due to low numbers of viable homozygous *Dicer2* mutant mosquitoes, an additional “mutant-enriched” G_3_ cross (6 homozygous mutant males mated with 3 homozygous mutant females, and 4 heterozygous females) was set up. The eggs from this cross were combined and hatched along with eggs from a pure heterozygous G_3_ cross (35 males and 33 females) to produce the G_4_ generation. Next, 7 males and 9 females homozygous for the *Dicer2* exon 5 mutation were crossed, establishing the homozygous mutant line. To ensure lack of contamination of the *Dicer2* mutant homozygous and control “sister” lines, at least 20 individuals from each generation have been routinely collected for PCR and Sanger sequencing surrounding the mutation site on exon 5 of the *Dicer2* gene. Throughout this article, the “sister line” is referred to as the control line, and the *Dicer2* mutant homozygous line is referred to as *Dicer2^null^* line.

### Mosquito exposure to infectious blood meals

Experimental infections of mosquitoes were performed in a biosafety level-3 containment facility, as previously described^55^. Shortly, 5- to 7-day-old female mosquitoes were deprived of 10% sucrose solution 24 hours before oral exposure to viruses. The infectious blood meal consisted of a 2:1 mix of washed human erythrocytes and viral suspension supplemented with 10 mM ATP (Sigma). The infectious titers were 10^7^ FFU/mL for DENV-1, 5 × 10^6^ FFU/mL for DENV-3, 10^6^ PFU/mL for ZIKV, 10^7^ PFU/mL for MAYV, and 10^6^ PFU/mL for CHIKV. Mosquitoes were offered the infectious blood meal for 15 min through a desalted pig-intestine membrane using an artificial feeder (Hemotek Ltd) set at 37°C. Fully engorged females were incubated at 28°C, 70% relative humidity and under a 12-hour light-dark cycle with permanent access to 10% sucrose.

### CFAV intrathoracic inoculation and RNA quantification

Mosquitoes were injected intrathoracically with 50 TCID_50_ units/mosquito of the CFAV-KPP strain using previously described methods^56^. Mosquitoes were collected on days 0, 2, 4, 6, 8 post inoculation and processed in 3 rounds of 4 samples per condition (inoculum × mosquito line × time point). Viral RNA was quantified by RT-qPCR as previously described^56^. CFAV RNA levels were normalized to the housekeeping gene *ribosomal protein 49* (*RP49*), and expressed as 2^−dCt^, with dCt = Ct^CFAV^ – Ct^RP49^.

### DENV RNA quantification

Mosquito body parts and organs were dissected in 1× PBS, and immediately transferred to a tube containing 800 μL of Trizol (Life Technologies) and ~20 1-mm glass beads (BioSpec). Samples were homogenized for 30 sec at 6,000 rpm in a Precellys 24 grinder (Bertin Technologies). RNA was extracted as previously described^57^, and stored at −80°C till further use. Viral RNA was reverse transcribed and quantified using a TaqMan-based qPCR assay, using DENV NS5-specific primers and 6-FAM/BHQ-1 double-labeled probe (sequences provided in Supplementary Table 1). Reactions were performed with the GoTaq® Probe 1-Step RT-qPCR System (Promega) following the manufacturer’s instructions. The limit of detection of the assay was 10 copies of viral RNA per μL.

### Virus titration

For CHIKV, MAYV and ZIKV, infectious titration was performed on confluent Vero cells plated in 24-well plates, 24 hours before inoculation. Ten-fold dilutions were prepared in DMEM alone and transferred onto Vero cells. After allowing infection, DMEM with 2% FCS, 1% P/S and 0.8% agarose was added to each well. Cells were fixed with 4% formalin (Sigma) after two (MAYV), three (CHIKV) or seven (ZIKV) days of incubation and plaques were manually counted after staining with 0.2% crystal violet (Sigma). DENV infectious titers were measured by standard focus-forming assay (FFA) in C6/36 cells as previously described^55^.

### Salivation assays

Mosquitoes were anesthetized on ice and wings and legs were removed from each individual. Proboscis was inserted into a 20-μL tip containing 20 μL of FBS for 30 min at room temperature (20-25°C). Saliva-containing FBS was expelled into 90 μL of Leibovitz’s L-15 medium (Gibco) for amplification and titration. The other tissues of interest (midgut, carcass, head) were dissected after salivations. Mosquito saliva samples were amplified in C6/36 cells for 5 days, and virus presence in amplified supernatants was assessed by virus titration.

### Survival assay

Mosquito were kept in 1-pint carton boxes with permanent access to 10% sucrose solution at 28°C and 70% humidity and mortality was scored daily. Experiments were performed in 3 to 4 replicate boxes of 21 to 38 mosquitoes each.

### Mosquito weighing

Five-to-seven-day old male and female mosquitoes were collected and starved for 24 hours prior to weighing to ensure that no sugar water remained in their digestive tracts. Mosquitoes were weighed in groups of 5 individuals of a single sex on a precision scale. Ten replicates were performed. Results are expressed as mass per individual mosquito.

### Pupation rates

Mosquito eggs were hatched for 30 min in a SpeedVac vacuum device (ThermoFisher). Two hundred larvae were counted and transferred to a standard rearing tray containing 1 L of tap water and a standard dose of fish food. Three to four separate trays were prepared for each mosquito line (control and *Dicer2* mutant). After 4-5 days of larval development, a daily count of the number of pupae (live and dead) was performed. The percentage of live and dead pupae was calculated relative to the total number of pupae recorded.

### Double-stranded RNA synthesis

Design and synthesis of dsRNA used in RNAi reporter assays has been described previously^57^. Briefly, dsRNA was synthesized from a GFP- or luciferase-containing plasmid. T7 promotor sequences were incorporated by PCR to the amplicon that was used as a template for the synthesis using the MEGAscript RNAi kit (Life Technologies). Primer sequences are provided in Supplementary Table 1.

### RNAi reporter assay

RNAi activity in adult mosquitoes was tested using a reporter assay described previously^33^. In short, 5 to 7-day-old adult female mosquitoes were intrathoracically injected using a Nanoject III (Drummond) with a suspension of ~200 nL containing a 1:1 mixture of unsupplemented Leibovitz’s L-15 medium (Life technologies) and Cellfectin II (Thermo Fisher Scientific) complexed with 50 ng pUb-GL3 (encoding Firefly luciferase, FLuc), 50 ng pCMV-RLuc (encoding Renilla luciferase), and 500 ng FLuc- or 1 μg GFP-specific specific dsRNA. After incubation for 3 days at 28°C, mosquitoes were homogenized in passive lysis buffer (Promega) using the Precellys 24 grinder (Bertin Technologies) for 30 sec at 6,000 rpm. Samples were transferred to a 96-well plate and centrifugated for 5 min at 12,000 × g. Fifty μL of supernatant were transferred to a new plate, and 50 μL LARII reagent added for the first FLuc measurement. Next, 50 μL Stop&Glow reagent was added before the second measurement of RLuc, according to the Dual Luciferase assay reporter system (Promega). Counts of RLuc were used to control for transfection efficiency, and samples with less than 1,000 counts were discarded from further analysis. Data were normalized by calculating the Fluc/RLuc ratio.

### Small RNA library preparation and sequencing

Total RNA from pools of 5 to 8 mosquitoes was isolated with TRIzol (Invitrogen). Small RNAs of 19-33 nucleotides in length were purified from a 15% acrylamide/bisacrylamide (37.5:1), 7 M urea gel as described previously^58^. Purified RNAs were used for library preparation using the NEB Next Multiplex Small RNA Library Prep kit for Illumina (E7300 L) with Universal miRNA Cloning Linker from Biolabs (S1315S) as the 3’ adaptor and in-house designed indexed primers. Libraries were diluted to 4 nM and sequenced using an Illumina NextSeq 500 High Output kit v2 (75 cycles) on an Illumina NextSeq 500 platform. Sequence reads were analyzed with in-house Perl scripts.

### Bioinformatics analysis of small RNA libraries

The quality of fastq files was assessed using graphs generated by ‘FastQC’ (http://www.bioinformatics.babraham.ac.uk/projects/fastqc/). Quality and adaptors were trimmed from each read using ‘cutadapt’ (https://cutadapt.readthedocs.io/en/stable/). Only reads with a phred score ≥20 were retained. A second set of graphs was generated by ‘FastQC’ on the fastq files created by ‘cutadapt’^59^. Mapping was performed by ‘Bowtie1’^60^ with the ‘-v 1’ option (one mismatch between the read and its target) using the following reference sequences: GenBank accession number HG316481 for DENV-1, GenBank accession number LN898104.1 for CHIKV, and positions 161,180,000-161,236,000 of chromosome 3 in the AaegL5 genome built^61^ for the histone transcript cluster. ‘Bowtie1’ generates results in ‘sam’ format. All ‘sam’ files were analyzed by different tools of the package ‘samtools’^62^ to produce ‘bam’ indexed files. To analyze these ‘bam’ files, different kind of graphs were generated using home-made R scripts with several Bioconductor libraries such as ‘Rsamtools’ and ‘Shortreads’ (http://bioconductor.org).

### Statistical analysis

Infection prevalence was analyzed as a binary variable by logistic regression and non-zero viral loads (infectious titers and RNA concentrations) were compared by analysis of variance (ANOVA) after log_10_ transformation. Each primary variable was analyzed with a full-factorial model including interactions up to the 2^nd^ order. Effects were considered statistically significant when *p* < 5%. The full test statistics are provided in Supplementary Figure 1 and Supplementary Figure 2. Because most of experiments were repeated multiple times, uncontrolled variation between experiments was accounted for in the statistical analyses. For graphical representations of viral loads, the data from different experiments were displayed in different colors. To show the statistical significance of pairwise comparisons the figures, a Student’s t-test or Mann-Whitney test was applied to the raw values after correction of the variation between experiments. This correction transforms the raw values into their deviation from the experimental mean, and the resulting values are centered around zero on the original scale. For the RNAi reporter assay, the graphical representation used the mean-centered values (after correction of the experiment effect) of the log_10_-transformed Firefly luciferase and Renilla count ratios. Estimates of transmission efficiency were compared with Fisher’s exact test. Pupation rates over time were analyzed with a 4-parameter model. The comparison of survival curves was performed using a Gehan-Breslow-Wilcoxon test. All statistical analyses were performed in JMP v.14.0.0 (http://www.jmp.com) or Prism v.9.3.1 (www.graphpad.com).

## Supplementary figures

**Supplementary Figure 1.**
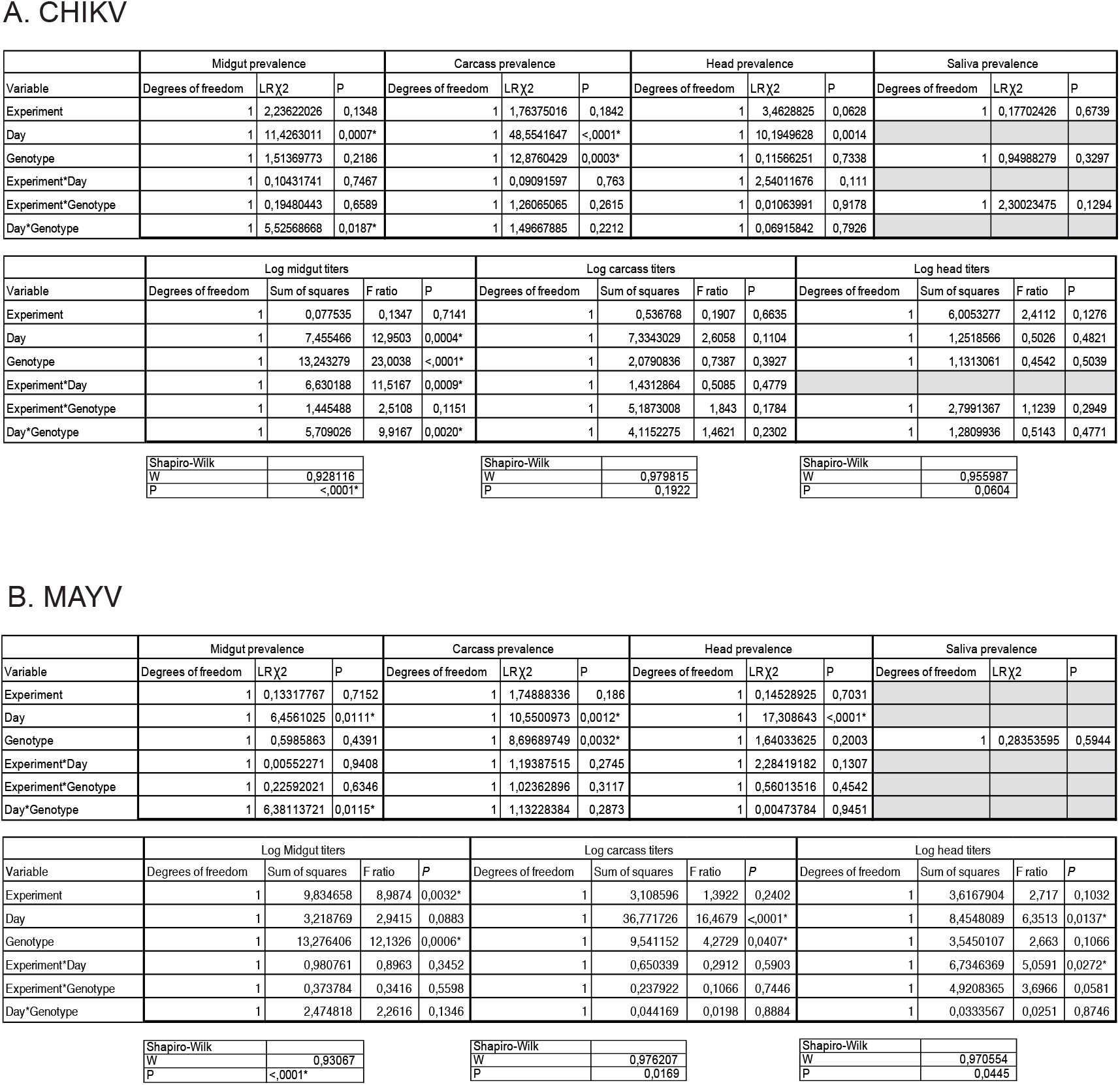
Detailed results of the statistical analysis of (**A**) CHIKV and (**B**) MAYV infection dynamics (Figure 3).

**Supplementary Figure 2.**
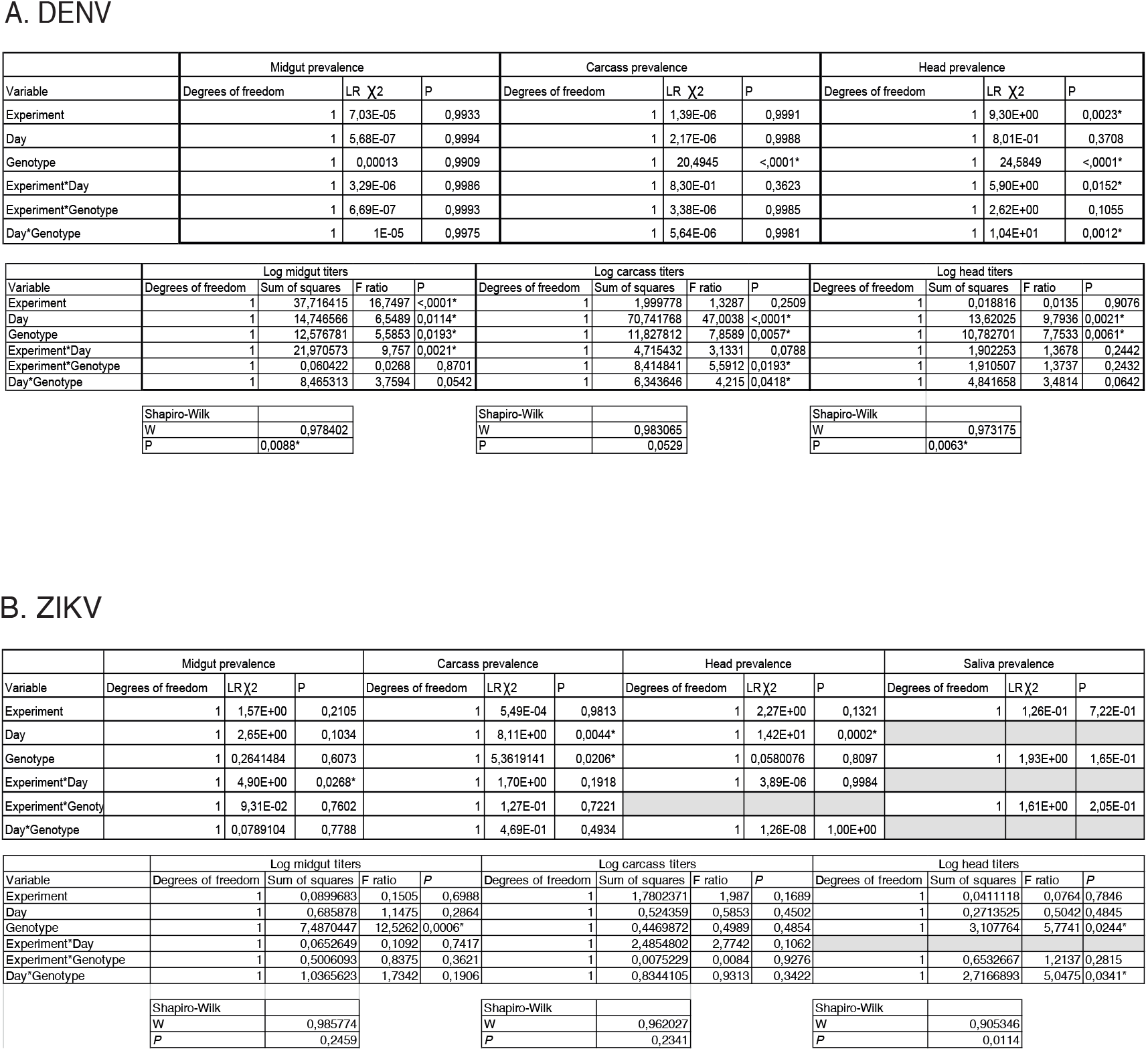
Detailed results of the statistical analysis of (**A**) DENV-3 and (**B**) ZIKV infection dynamics (Figure 4).

**Supplementary Figure 3.**
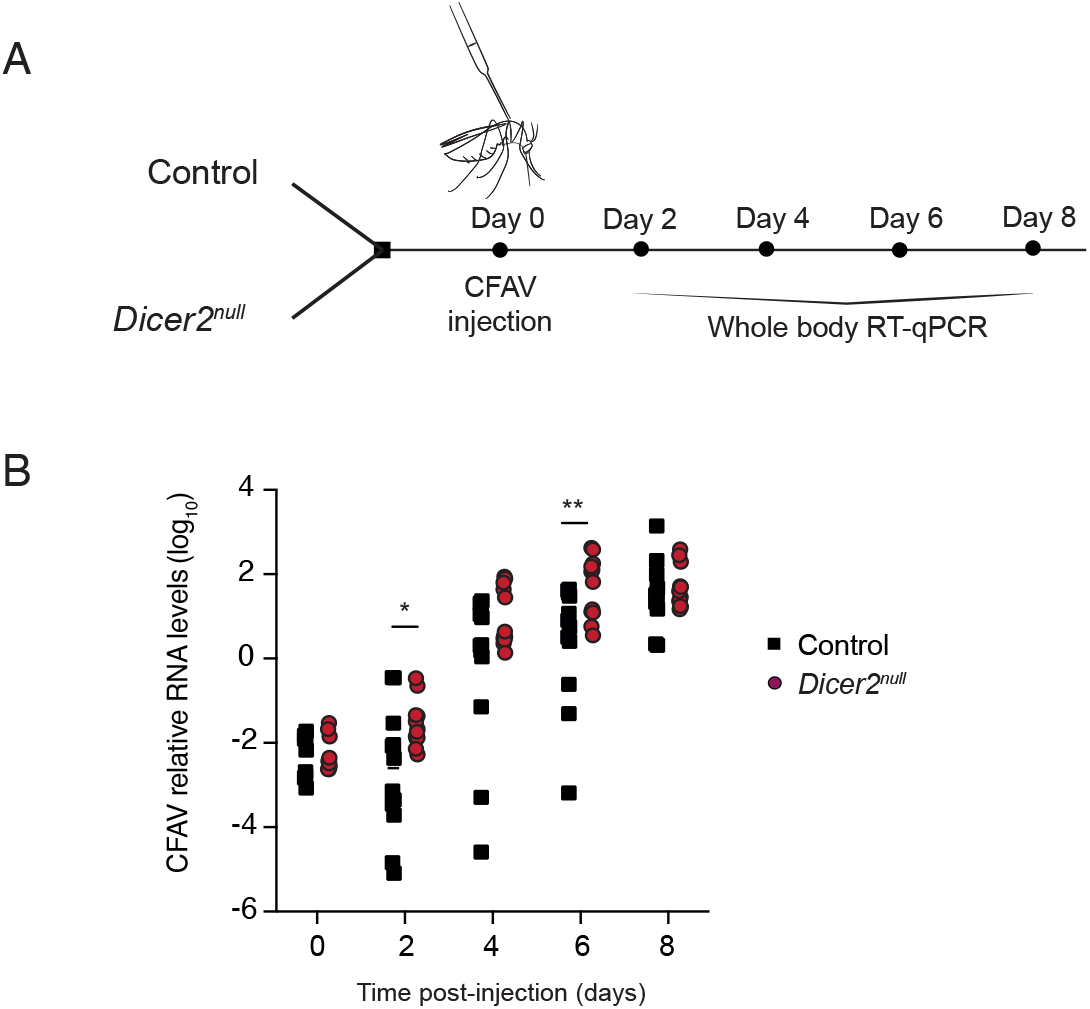
(**A**) Experimental scheme for the analysis of CFAV infection dynamics. Adult *Dicer2^null^* mutants and control female mosquitoes were inoculated intrathoracically with 50 TCID_50_ units of CFAV. Whole mosquito bodies were collected on days 0, 2, 4, 6 and 8 post injection and relative viral RNA levels were measured by RT-qPCR. (**B**) Relative CFAV RNA levels over time post inoculation in control and *Dicer2^null^* mutant mosquitoes. Each experimental condition is represented by 9-12 mosquitoes. Statistical significance of the pairwise differences was tested with a Mann-Whitney test (*p<0.05; **p<0.01; ***p<0.001).

**Supplementary Figure 4.**
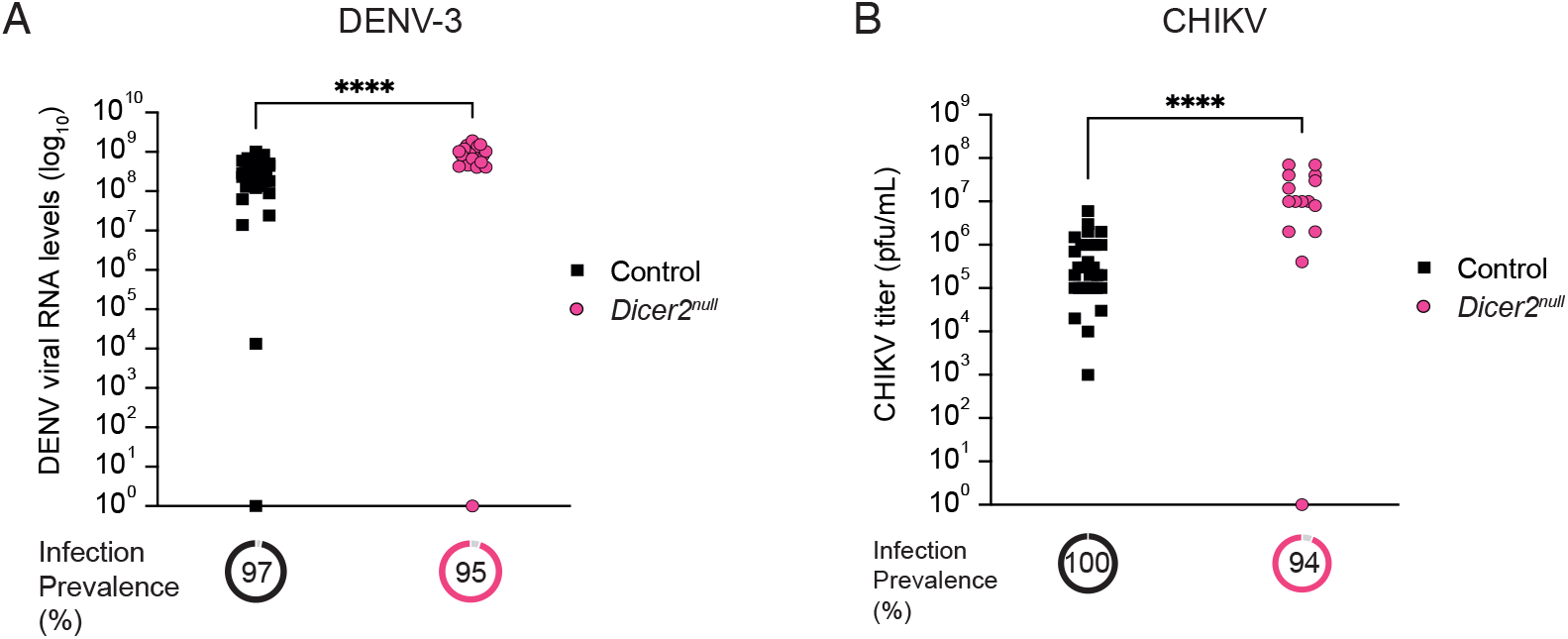
Infection levels in mosquitoes of the survival analysis shown in Figure 6. Infection prevalence in *Dicer2^null^* mutants and control female mosquitoes was measured 7 days after exposure to a (**A**) DENV-3 or (**B**) CHIKV infectious blood meal by (**A**) RT-qPCR or (**B**) viral titration. Each experimental condition is represented by 15-32 mosquitoes. Statistical significance of the pairwise differences was tested with a Mann-Whitney test (*p<0.05; **p<0.01; ***p<0.001).

**Supplementary Table 1.**
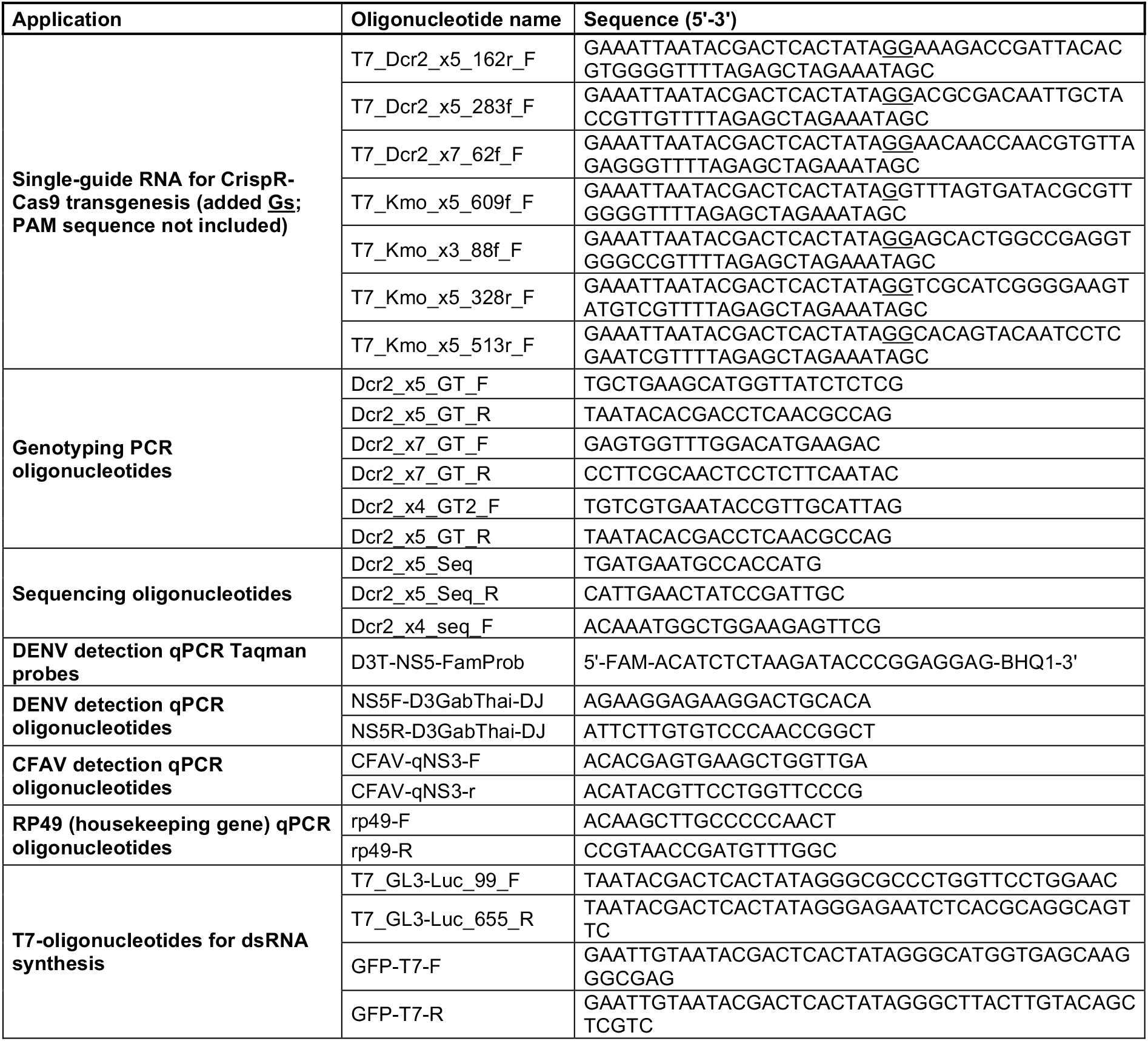
List of oligonucleotides.

